# Significant antitumour effect of an antibody against TMEM180, a new cancer-specific molecule

**DOI:** 10.1101/397844

**Authors:** Masahiro Yasunaga, Shinji Saijou, Shingo Hanaoka, Takahiro Anzai, Ryo Tsumura, Yasuhiro Matsumura

## Abstract

The present state of therapy for colorectal cancer (CRC) is far from satisfactory, highlighting the need for new targets for this disease. We identified a new colorectal cancer (CRC)-specific molecule, TMEM180, a predicted eleven-pass transmembrane protein that apparently functions as a cation symporter. Our anti-TMEM180 monoclonal antibody (mAb) eradicated SW480 CRC xenografts in mice. The *TMEM180* promoter region contains ten hypoxia-responsive element consensus sequences; accordingly SW480 cells upregulated TMEM180 under low-oxygen conditions. TMEM180 expression in SW480 cells was positively correlated with anchorage-independent colony formation and tumourigenesis. TMEM180-positive SW480 cells resided at the tumour–stroma interface manifested by αSMA-positive fibroblasts, also known as the tumour niche. Some clusters of TMEM180-positive cells adjacent to the niche were integrin α6-positive. These data indicate that TMEM180 represents a possible cancer stem cell marker and that a mAb against this protein could be used as antibody-based therapeutic against CRC.

## Introduction

Colorectal cancer (CRC) is the third leading cause of cancer-related mortality in the world (WHO, 2002). Despite recent progress in chemotherapeutic options, including monoclonal antibody (mAb) therapeutics, the prognosis for metastatic CRC is still very poor. Of the mAb therapeutics for CRC, mAbs against vascular endothelial growth factor (VEGF) and epidermal growth factor (EGFR) are clinically available^1,2^. Although EGFR is expressed in several normal tissues, the mAb can preferentially accumulate in CRC tumour tissues due to the enhanced permeability and retention effect of tumour tissue^3,4^. However, because EGFR is expressed at a high level in normal skin tissue, skin toxicity is common, and cessation of mAb treatment is sometimes inevitable even if the therapy is effective in patients^5,6^. In the case of anti-VEGF mAb, some patients receiving the mAb have a life-threatening side effects including serious bleeding and gastrointestinal perforation^7^.

In this context, we need to find a new CRC specific molecule and develop the mAb against the molecule in order to produce the drastic antibody therapeutics in the treatment of metastatic CRC with minimal side effects.

To find a CRC-specific molecule, a comprehensive expression analysis is usually performed between CRC cells and their normal counterpart, mucoepithelial cells. However, in reality, obtaining pure live normal mucoepithelial cells is more difficult than obtaining CRC cells even if laser micro-dissection method is used. We had reported that many mucoepithelial cells are exfoliated in stool, some of which contain intact mRNA^8^. Based on the results, we previously developed a method for obtaining almost pure normal mucoepithelial cells from the lavage solution following colonoscopies of healthy examinees. We then identified several new CRC specific molecules after the comprehensive expression analyses between the pure mucoepithelial cells and the CRC cell lines. TMEM180 is one of them and a predicted eleven-pass transmembrane protein (UniProtKB, http://www.uniprot.org/). In the present study, we developed a mAb against TMEM180 and clarified the CSC-like effects of TMEM180 and evaluated the potential of using its mAb for CRC therapy.

## Results

### Identification and characterization of TMEM180 as a new CRC marker

We previously developed a method to obtain almost pure normal mucoepithelial cells from the lavage solution following colonoscopies of healthy examinees using immune-beads with the anti-EpCAM mAb^9^. Moreover, a previous study reported a comprehensive DNA microarray analysis comparing CRC cell lines (SW480, LoVo, DLD-1, HT-29 and HCT116) and normal mucoepithelial cells^9^. We reanalyzed these microarray data and found that TMEM180 was highly expressed in five CRC cell lines but not in the normal colonocytes obtained from lavage of two healthy volunteers (Fig. 1a). In quantitative RT-PCR (qPCR) analysis, performed as a first validation, TMEM180 was expressed at high levels in CRC tissue samples (Fig. 1b). *In situ* hybridization (ISH) was then conducted for the final validation (Fig. 1c, d). All of these methods indicated that TMEM180 was a new CRC marker (Fig. 1a-d).

**Figure 1.**
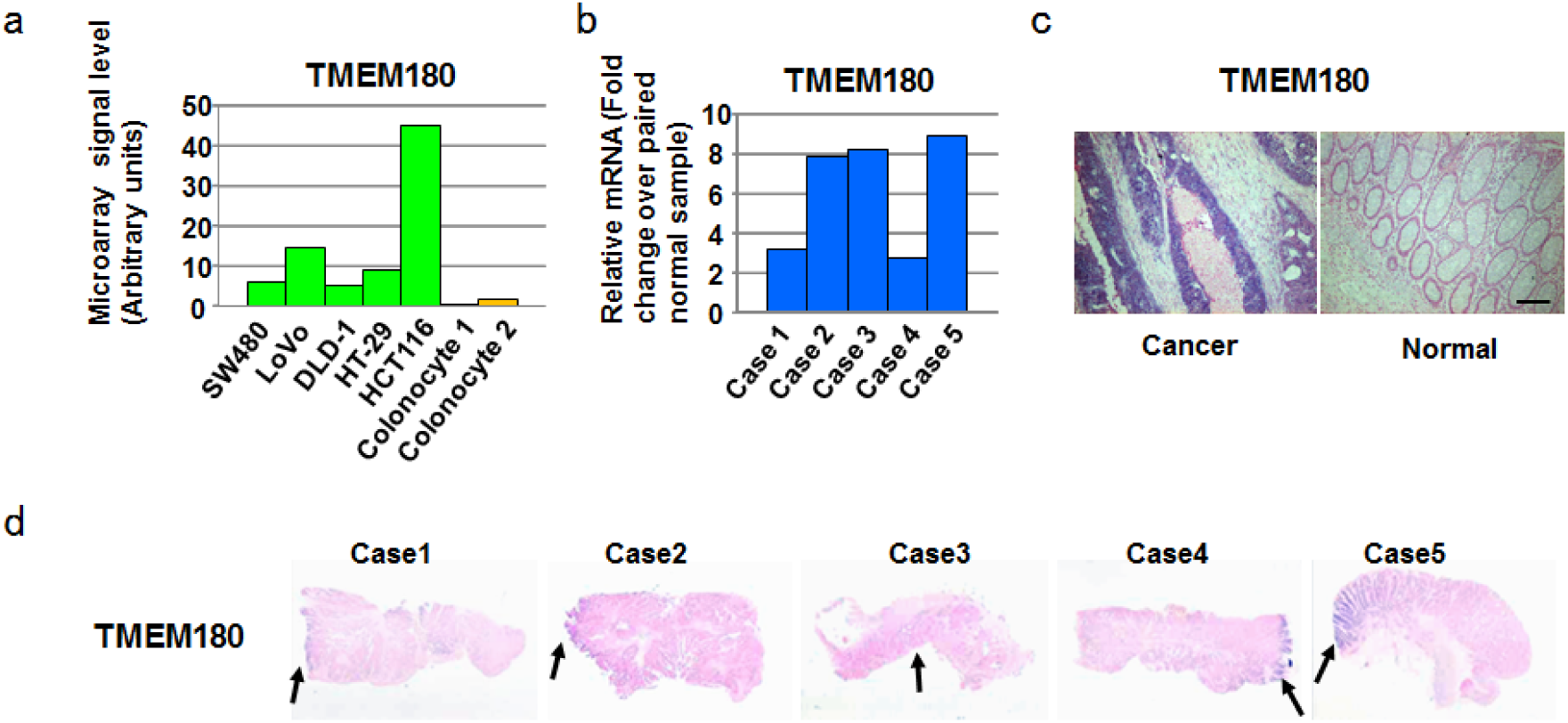
Identification of TMEM180 as a colorectal cancer-specific marker. a. *TMEM180* gene expression in five CRC cell lines and two colonocyte samples from healthy donors was evaluated using DNA microarray analysis. b. *TMEM180* gene expression in five clinical samples was evaluated using quantitative RT-PCR. Relative quantification as tumour-to-normal tissue ratio. c and d. ISH (*in situ* hybridization) of TMEM180 in the clinical samples. Arrows indicate cancer (d).

### Diagram and functional role of TMEM180

TMEM180 is classified in the major facilitator superfamily and appears to function as a cation symporter (Fig. 2a). To address the functional role of TMEM180, we established two *TMEM180* gene knockdown (KD) SW480 cell lines: KD1 cells with the lowest expression of TMEM180 protein, and KD2 cells with intermediate TMEM180 protein expression (Fig. 2b). We investigated the uptake of glutamine and arginine in the cation symport mechanism because the utilization of both amino acids was strongly increased in tumour cells ^10,11^. We found that SW480-WT cells were capable of growing in a serum-free medium with glutamine and arginine. By contrast, both KD1 and KD2 could not grow in the same medium (Fig. 2c). These findings suggest TMEM180 has an important role in the uptake or metabolism of glutamine and arginine in tumour growth and proliferation.

**Figure 2.**
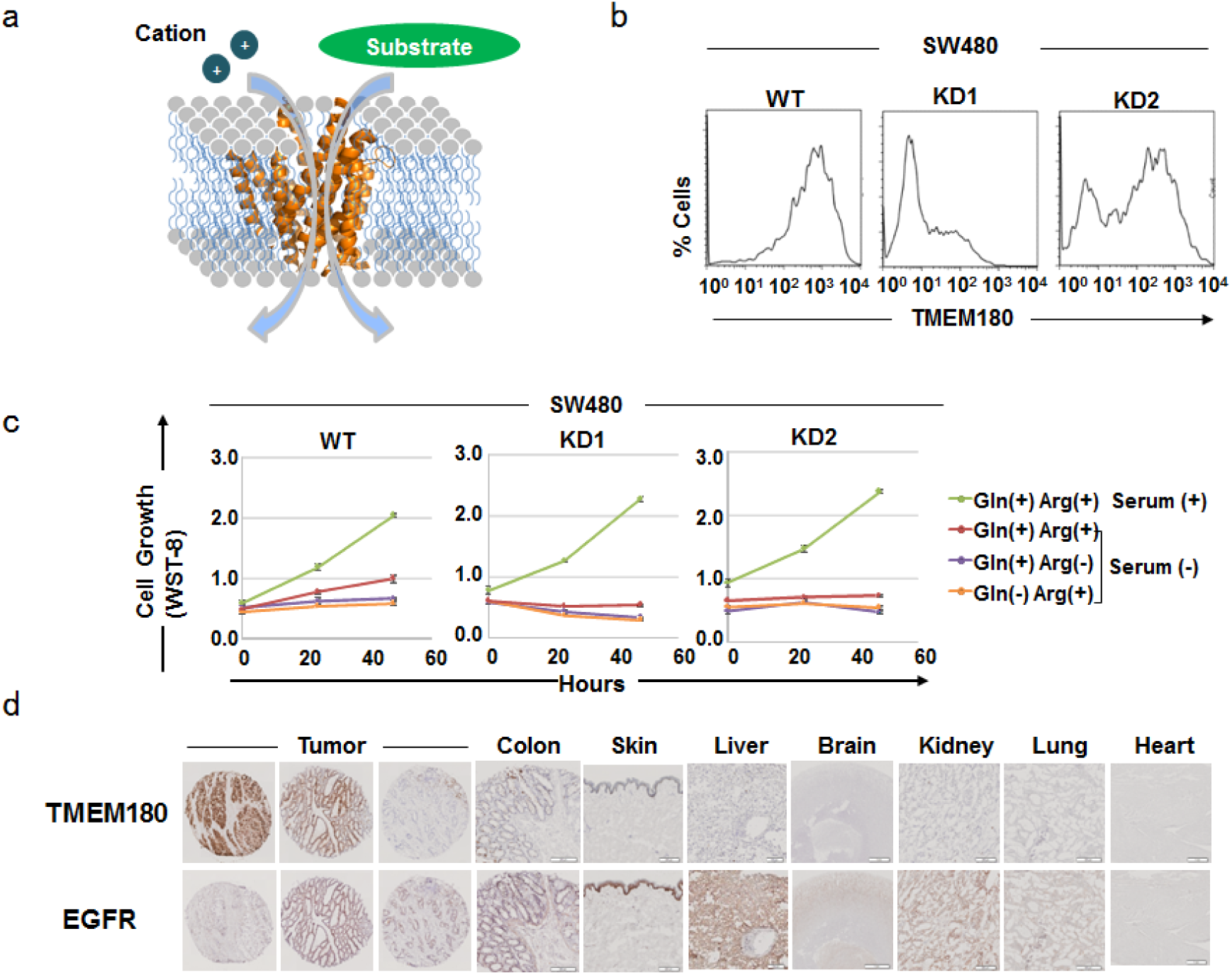
Diagram and functional role of TMEM180. a. Diagram of TMEM180 as a cation symporter is shown. b. Flow cytometric analysis of the SW480 wild-type (WT), gene knockdown clone No. 1 (KD1) or gene knockdown clone No. 2 (KD2) cells. The difference in the TMEM180 expression level in SW480-WT, -KD1 and -KD2 cells is shown. c. Cell growth of SW48-WT, -KD1 or -KD2 cells in (1) glutamine (Gln)–positive (+), arginine (Arg) (+) and Serum (+), (2) Gln (+), Arg (+) and Serum-negative (-), (3) Gln (+), Arg (-) and Serum (-) or (4) Gln (-), Arg (-) and Serum (-). Each growth activity was evaluated by WST-8 assay. d. Immunohistochemistry showing the expression of TMEM180 or EGFR in CRC and normal tissues (colon, brain, heart, lung, liver, kidney and skin). Scale bar = 100 μm.

### Release of TMEM180-positive exosomes from CRC cells

To evaluate the potential of TMEM180 as a new target for the diagnosis and/or therapy of CRC, we attempted to produce a mAb. However, the *TMEM180* gene is predicted to encode an eleven-pass transmembrane protein with limited extracellular domain expression. First, we immunized rats with a recombinant extracellular domain of the TMEM180 protein and obtained an IgM mAb. During the evaluation process, we unexpectedly noticed that the TMEM180 protein was present in the culture supernatant of SW480 and DLD-1 CRC cells. We collected the supernatant and subjected it to hydroxyapatite and gel filtration chromatography. We found that a high concentration of the TMEM180 protein existed in the void fraction with a molecular weight of over 600 kDa (Supplementary Fig. 1a).

Due to the detection of TMEM180 in culture supernatant, we hypothesised that TMEM180 coexisted with exosomes or other extracellular microparticles. To test whether TMEM180 exists on tumour-derived exosomes, we conducted immune-electron microscopy for the enriched exosomes with the CD9 or CD63 mAbs and the TMEM180 mAb. The images clearly indicated that TMEM180 was present on the tumour exosome, similar to CD9 and CD63 (Supplementary Fig. 1b). In the sandwich ELISA, both TMEM180-positive and CD9-positive exosomes were detected (Supplementary Fig. 1c). We concluded that the tumour exosomes released from DLD-1 cells possess TMEM180.

### Production of anti-TMEM180 antibody and validation of a CRC-specific target

Because we judged that purifying the full-length TMEM180 protein while maintaining the native structure necessary for producing a useful mAb recognizable for the live cell would be the hardest step, we immunized rats again with TMEM180-positive tumour exosomes purified from the supernatant of DLD-1 cells. The newly obtained anti-TMEM180 mAb (IgM, clone 669) was able to recognize most CRC cells, but not haematopoietic cells (Supplementary Fig. 2a, b).

We then converted the rat IgM to humanized IgG1 in order to evaluate the potential for therapeutic application of the mAb. First, we examined the expression of TMEM180 in CRC and normal major tissues by immunohistochemistry (IHC). Cetuximab (an anti-EGFR mAb) was used as a reference as it is currently widely used for CRC patients. We found that 9 of the 37 CRC tissues (24.3%) were TMEM180-strong positive and 43.2% (16/37) were TMEM180-weak positive. Regarding EGFR, 7 of 37 CRC tissues (18.9%) were EGFR-strong positive and 35.1% (13/37) were EGFR-weak positive. However, there was no significant difference in the expression level between TMEM180 and EGFR (Fig. 2d, Supplementary Fig. 3a, b). IHC with the mAb did not show staining in major organs including the brain, heart, lung, liver, kidney, colon and skin (Fig. 2d). In contrast, IHC with cetuximab showed strongly positive staining in the skin, moderately positive staining in the liver and colon and weakly positive staining in the brain, kidney and lung (Fig. 2d).

According to a database, the expression of TMEM180 at the mRNA level is very low not only in normal organs but also in their respective cancer tissues (Supplementary Fig. 4a, b). Our data indicate that TMEM180 is highly expressed in CRC.

### The anti-TMEM180 mAb is highly effective in the targeted inhibition of CRC tumours

The anti-TMEM180 mAb showed little to no direct cytotoxicity in DLD-1 and SW480 cells *in vitro* (Supplementary Fig. 5a). We next evaluated the antibody-dependent cell-mediated cytotoxicity (ADCC) and complement-dependent cytotoxicity (CDC) in DLD-1 and SW480 cells, both of which are Ras mutation-positive. Cetuximab was also used in this test as a reference. The anti-TMEM180 mAb appeared to possess ADCC and CDC in both cell lines (Supplementary Fig. 5b, c).

Because of *in vitro* ADCC activity of both mAbs, we evaluated the *in vivo* anti-tumour activity of the anti-TMEM180 mAb and cetuximab in mice bearing DLD-1 or SW480 xenografts. Preliminary tests showed that ceuximab exerted statistically significant antitumour effects in both tumour xenografts. In the case of anti-TMEM180 mAb, the antitumour effect was significant in DLD-1 xenografts. Strikingly, all SW480 xenografts were completely eradicated by the anti-TMEM180 mAb. We then confirmed these results using a dosage of 25 or 12.5 mg/kg given twice a week for a total of eight injections for both mAbs in mice bearing DLD-1 or SW480 xenografts, respectively. In mice with the DLD-1 xenografts, both anti-TMEM180 mAb and cetuximab showed a significant antitumour effect compared to the control treatment (Fig. 3a). In mice with the SW480 xenografts, cetuximab showed a significant antitumour effect at 25 mg/kg but not at 12.5 mg/kg. On the other hand, anti-TMEM180 mAb elicited a striking antitumour effect, eradicating the tumour at both 25 and 12.5 mg/kg (Fig. 3a). There was no significant difference in the treatment-related body weight loss among the five groups (Fig. 3b).

**Figure 3.**
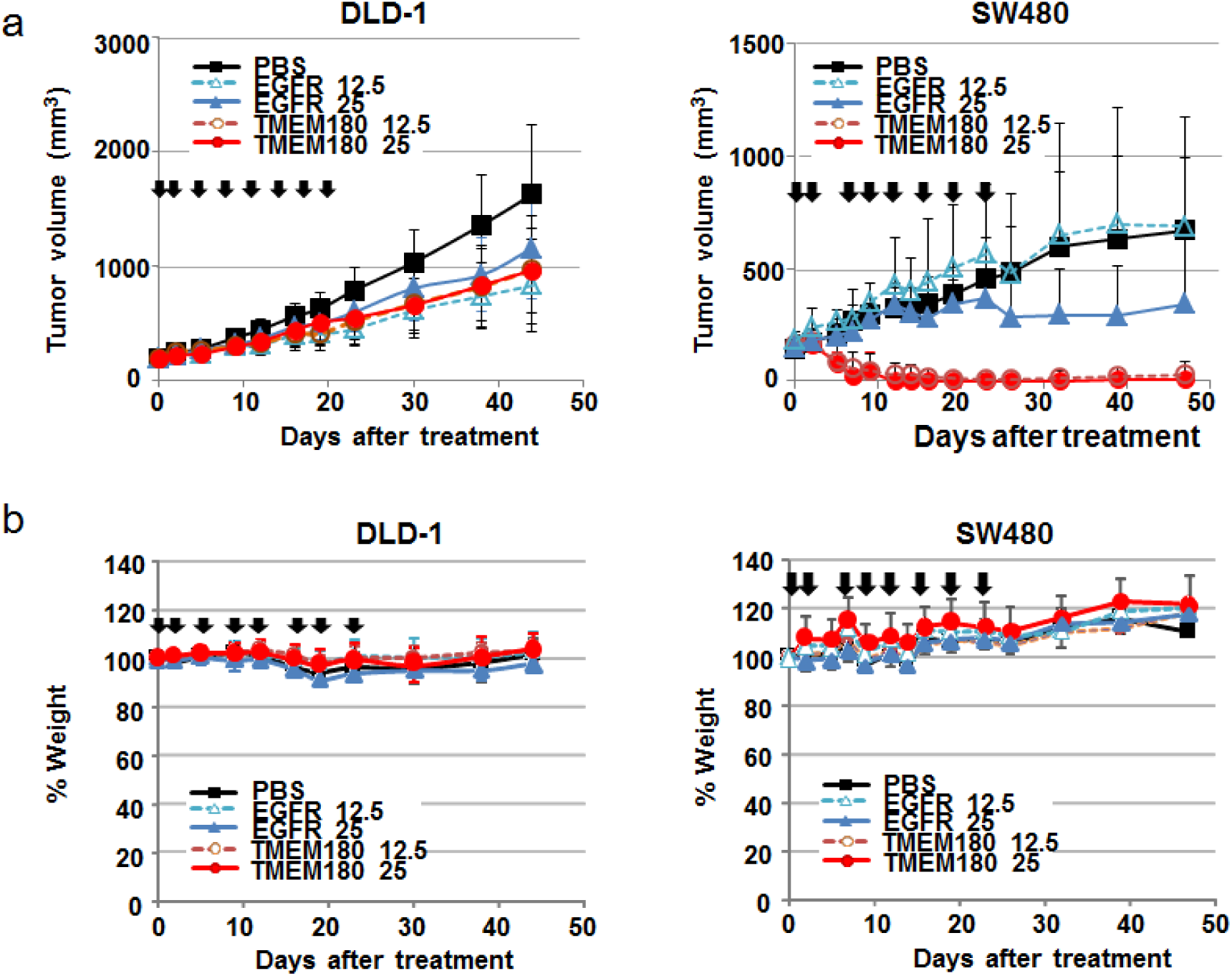
Anti-tumour effect of the anti-TMEM180 antibody against CRC tumours. a and b. Anti-tumour activity of the anti-TMEM180 mAb against DLD-1 (a) and SW480 (b) xenograft models. When the tumour volume reached approximately 200 mm^3^, each treatment started (day 0). The anti-TMEM180 mAb (=TMEM180), the anti-EGFR mAb cetuximab (=EGFR) or saline (=PBS) as a control was administered at a dose of 12.5 or 25 mg/kg to each group of mice (n = 5) by intraperitoneal injection twice a week from Day 0 to 22. The arrows indicate the days of administration, and the curves show the effect of the treatment on tumour size. P < 0.001 (SW480 xenograft, PBS compared with TMEM180 12.5 mg/kg or 25 mg/kg; EGFR 12.5 mg/kg compared with TMEM180 25 mg/kg), P < 0.01 (DLD-1 xenograft, PBS compared with EGFR 25 mg/kg, TMEM180 12.5 mg/kg or 25 mg/kg; SW480 xenograft, EGFR 25 mg/kg compared with TMEM180 12.5 mg/kg). P < 0.05 (DLD-1 xenograft, PBS compared with EGFR 12.5 mg/kg). Bar = SD. c and d. Percent change in body weight in the same mice with the same treatments as in a and b. Bar = SD.

The question is why such striking antitumour effect by the anti-TMEM180 mAb was obtained only in the SW480 xenografts because both DLD-1 and SW480 cells showed equal TMEM180 expression. This eradication of SW480 xenografts by the anti-TMEM180 mAb suggested the existence of CSCs in the tumours and these results prompted us to conduct following intensive experiments connected to characteristics of CSCs.

### TMEM180 expression is up-regulated by hypoxia in CRC cells

Using a public database to analyse the promoter region of the *TMEM180* gene, we found that there were ten hypoxia response element consensus sequences^12,13^ in the gene (Fig. 4a). Moreover, HIF-1α expression was elevated in both DLD-1 and SW480 cells in hypoxic conditions (Fig. 4b). We therefore evaluated the change in TMEM180 expression in response to the same treatment. Hypoxia significantly increased *TMEM180* mRNA expression in SW480 cells, and also tended to increase expression in DLD-1 cells (Fig. 4c). The protein expression level of TMEM180 was significantly increased in both DLD-1 and SW480 cells after hypoxia treatment (Fig. 4d, e). Therefore, TMEM180 expression in CRC cells is regulated by hypoxia.

**Figure 4.**
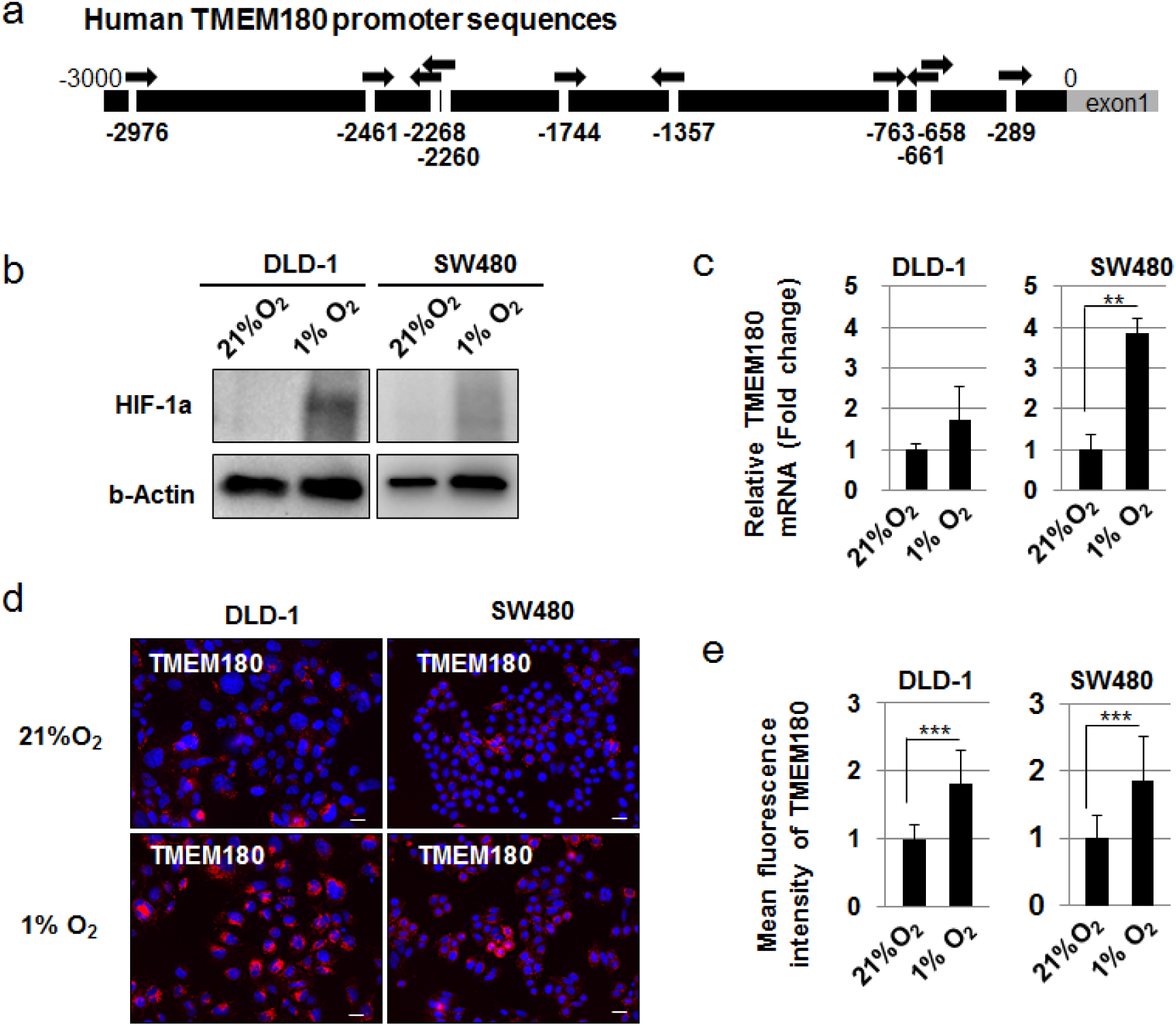
TMEM180 expression is controlled by hypoxia in CRC cells. a. Schema of the promoter in the *TMEM180* gene. Ten hypoxia response element consensus sequences (HREs) were found in the promoter sequence. HREs are indicated by arrows. b. Western blotting of HIF-1α. The difference in the HIF-1α protein expression level in the colorectal cancer (CRC) cell lines DLD-1 and SW480 under normoxia (21% O_2_) and hypoxia (1% O_2_) is shown. The expression of β-Actin was used as a control. c. Quantitative RT-PCR of *TMEM180*. The difference in the *TMEM180* mRNA expression level in DLD-1 and SW480 cells under normoxia (21% O_2_) and hypoxia (1% O_2_) is shown. **P < 0.01. Bar = SD. d and e. Immunostaining of TMEM180. The change in the TMEM180 expression pattern in DLD-1 and SW480 cells under normoxia (21% O_2_) and hypoxia (1% O_2_) is shown (D). Scale bar = 10 μm. The data were quantified and are shown in (E). ***P < 0.001. Bar = SD.

### TMEM180 has an important role in maintaining CSC properties

The CSC hypothesis suggests that tumour cells are heterogeneous and that only the CSC subpopulation has the ability to maintain the capacity to self-renew, proliferate extensively and form new tumours^14-19^. The CSC niche, which is a specific anatomical location within the tumour microenvironment, is necessary to maintain the principle properties of CSCs^20,21^. A hypoxic microenvironment is well known to be important for CSC niche formation^22-24^. Very recently it is reported that sub-populations of CSCs located at the tumour-stroma interface are rich in integrins^25,26^. Therefore, we evaluated the localization of TMEM180-positive CRC cells involved in niche formation. We found that TMEM180-positive tumour cells closely localized with the αSMA-positive tumour microenvironment as a hypothetical niche in SW480 tumour xenografts and some clusters of TMEM180 positive cells close to the niche were integrin α6-positive (Fig. 5a). Importantly, SW480 tumour xenografts were completely eradicated, even if SW480 xenografts contain extensive area of TMEM180-negative region (Fig. 5a). By contrast, the DLD-1 xenografts did not possess a clear αSMA-positive tumour microenvironment, although a homogeneous distribution of TMEM180-positive tumour cells was observed (Fig. 5a).

**Figure 5.**
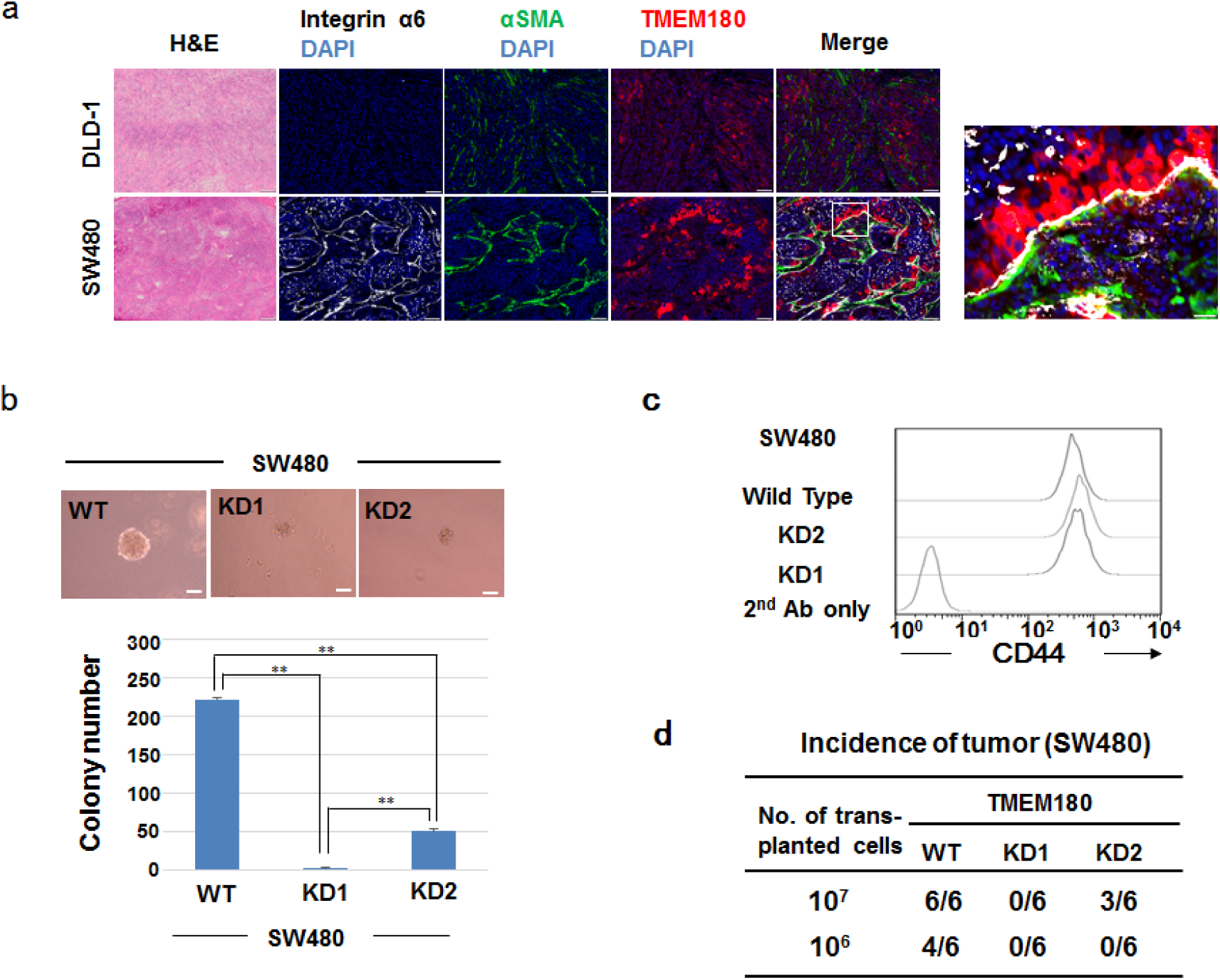
TMEM180 plays in CSC properties in CRC cells. a.Immunostaining of TMEM180-positive CRC cells with or without the tumour niche. αSMA (green), TMEM180 (red), Integrin α6 (white) and nuclei (blue) of a DLD-1 tumour and SW480 tumour are shown. Scale bar = 100 μm. Right, higher magnification of boxes area. Scale bar = 20 μm. b. *In vitro* colony-forming activity of SW480-WT, -KD1 and -KD2 cells. Representative images are shown (left). Scale bar = 50 μm. The number of colonies greater than 50 μm in diameter was counted (right). **P < 0.01. Bar = SD. c. CSC marker CD44 expression in SW480-WT and –KD1 and –kD2 cells. The representative expression level of CD44 using flow cytometry was shown. d. *In vivo* tumour initiation activity of SW480-WT, -KD1 and -KD2 cells was measured by evaluating the tumour incidence at the implanted sites on BALB/c nude mice 6 weeks after injection.

Next, we examined whether TMEM180 affected the CSC properties in SW480 cells. In an assay examining *in vitro* colony-forming activity in soft agar, SW480-WT cells clearly formed colonies, but neither KD1 nor KD2 cells did (Fig. 5b). By contrast, CSC marker CD44 being well-used for the characterization ^27-29^, was not changed among controls, KD1 and KD2 cells (Fig. 5c). On the other hand, the *in vivo* tumour-initiating activity of the SW480 cell lines was positively correlated to the level of TMEM180 expression (Fig. 5d). Although CD44 was not changed, both in vitro colony-forming activity and in vivo tumour-initiating activity as the principle CSC properties were lost in correlation with the loss of TMEM180 expression.

## Discussion

TMEM180 mAb staining was almost absent in normal tissues, unlike staining with the EGFR mAb. Anti-TMEM180 mAb treatment suppressed DLD-1-tumour growth equally to cetuximab treatment. Surprisingly, SW480 tumours were completely eradicated by anti-TMEM180 mAb treatment but not by cetuximab treatment.

In further experiments, we found that the *TMEM180* gene had ten hypoxia response element consensus sequences, which may be HIF-1α binding sites controlled by hypoxia^12,13,30^. In the SW480 xenografts, TMEM180-positive SW480 tumour cells resided close to the αSMA-positive stroma area, the so-called tumour niche, which is indispensable for maintaining the stemness of CSCs^31,32^. Moreover, hypoxia is strongly associated with tumour niche formation^24,33,34^. TMEM180-positive SW480 cells were developed in the tumour hypoxic niche to become robust CSCs. Recent studies have indicated that not only hypoxia but also chemotherapy induces HIF-1α expression followed by T-cell anergy and increased dissemination of cancer cells^35^. HIF-1α is currently considered a key factor of CSC function. In cancer tissues, unlike normal organs, angiogenesis and hypoxia associated with the necrosis caused by cancer-induced blood coagulation are repeated alternatively during the progression in clinic^36-38^. Further basic and translational studies are needed to elucidate the detailed mechanism by which TMEM180 signalling modulates the CSC properties in association with the tumour hypoxic niche. In addition, we found that integrin 6α was clearly expressed in some clusters of TMEM180-positive cells adjacent to the niche, which was consistent with CSC criteria reported previously^25,26^. However, we cannot conclude whether TMEM180 is a new CSC marker or not because the molecule does not correspond to all criteria of CSC such as CD44 expression state. We need to conduct additional studies to make clear this issue.

The safety of a drug should also be considered for therapeutic use in clinical practice. According to the database, *TMEM180* mRNA is hardly expressed in any normal tissues (Supplementary Fig. 4a). Also, the expression of TMEM180 is extremely low in all organs compared to the expression of other known CRC markers (Supplementary Fig. 6). The IHC in this study also showed that there was no TMEM180 expression in all major organs studied. Finally, we succeeded in obtaining a *TMEM180* gene knockout mouse that does not exhibit embryonic, neonatal, and postnatal lethality (Supplementary Table 1). According to these data, we expect that unpredictable adverse effects are unlikely to occur in a clinical trial of the anti-TMEM180 mAb. TMEM180 was expressed at levels as high as the levels of other existing markers such as EGFR, CD44 and EpCAM in cancers and at extremely low levels compared to the levels of other molecules in normal tissues. We, therefore, think that the tumour specificity of TMEM180 is substantially higher than that of other molecules. Our mAb is now being processed according to Good Manufacturer Practice principles for testing in a first-in-human clinical trial.

However, we think that there are several issues to be clarified before entering clinical trial. We have to investigate the further antitumour activity using other CRC cell lines and PDX tumours. We will make progress the functional analysis of TMEM180 after obtaining F2 generation of the KO mice. We hope that future clinical trial will verify the significance of the mAb for the treatment of CRC.

## Online Methods

### Collection of human samples

Human exfoliated colonocytes in a saline mucosal wash fluid were obtained from two healthy donors at the time of their colonoscopy examination. The cell purification and processing methods were conducted as previously described^9^.

The experimental protocols and procedures were approved by the institutional review board of the National Cancer Center, Japan. All methods were performed in accordance with the relevant guidelines and regulations. Two healthy volunteers in this study provided written informed consent.

### Cells and cell culture

The human colon cancer cell lines of SW480, LoVo, DLD-1, HT-29, HCT116, WiDr, and Colo320 and haematopoietic cell lines of K562, Ramos, RL and Raji were purchased from the American Type Culture Collection. TMEM180 gene knockdown SW480 cells were established according to our previously reported protocol^39^. For the overexpression of TMEM180 gene, plasmid pCMV-TMEM180 (Origene) was transfected into DLD-1 cells (DLD-1-OE) using Lipofectamine LTX (Thermo Fisher Scientific) according to the manufacture’s protocol.

The growth and survival of the cells were evaluated using the WST-8 assay (Cell Counting Kit-8, Dojindo). The reaction signals were evaluated by measuring the absorbance at 450 nm and using a microplate reader (SpectraMax Paradigm, Molecular Devices Corp.).

To evaluate the cellular response to hypoxia, DLD-1 and SW480 cells were cultured in DMEM (Wako) supplemented with 10% FBS (Life Technologies) and a 1% penicillin-streptomycin-amphotericin B suspension (Wako) under normoxic (37°C, 5% CO_2_, 21% O_2_) or hypoxic (37°C, 5% CO_2_, 1% O_2_) conditions. The BIONIX2 hypoxic cell culture kit (Sugiyamagen) was also used to generate hypoxic conditions according to the manufacturer’s instructions.

To evaluate the colony-forming activities, the bottom layer composed of 1 volume of 1.8% agar solution (Lonza) and 1.6 volumes of the cultured medium with 0.15% NaHCO_3_ was prepared. The cells were suspended in the top mix layer composed of 1 volume of 1.8% agar solution (Lonza) and 4 volumes of the cultured medium with 0.2% NaHCO_3_. A single cell suspension obtained using a 40-μm cell strainer (Corning Incorporated) at a final concentration of 2 × 10^5^ cells/ml was poured on the bottom layer. After 1 or 2 weeks, the number of colonies with a size over 50 μm was counted.

### DNA microarray

DNA microarray analysis was conducted using 10 µg of total RNA from colon cancer cells or exfoliated colonocytes and the Affymetrix GeneChip Human Genome U133 plus 2.0 arrays; the analysis was conducted according to the standard Affymetrix protocols^40^. In the microarray analysis, 91 genes encoding membrane proteins, which were highly expressed in CRC cells but not in the colonocytes from healthy donors, were selected from 38,500 genes contained in the chip array.

### Real-time quantitative RT-PCR

RNA extraction and qRT-PCR were conducted as previously described^41^. The PCR reaction consisted of 10 µl TaqMan Fast Universal PCR Master Mix (Thermo Fisher Scientific), 1 µl TaqMan primer/probe mixture (Thermo Fisher Scientific) and 9 µl template cDNA diluted to a total volume of 20 µl. Real-time PCR was performed in an Applied Biosystems 7500 Fast System (Thermo Fisher Scientific). The relative quantification of the total RNA in each sample was conducted using the comparative Ct (threshold cycle) method. The relative expression of each gene was normalized against expression of *GAPDH* or *ACTB*.

### *In situ* hybridization

*In situ* hybridization was conducted as previously described^9^.

The deparaffinized sections of human tissue (Genostaff Co., Ltd.) were fixed with 4% paraformaldehyde (PFA, Wako). After treatment with 7 μg/mL proteinase K (Roche), the sections were refixed with 4% PFA and acetylated with 0.25% acetic anhydride. After the tissue was dehydrated, hybridization was performed with digoxigenin-labelled RNA probes (475-bp fragment of TMEM180, gene bank accession number NM_024789, nucleotide positions 1314–1789). After the tissue was washed, RNase treatment was conducted. The sections were then rewashed and treated with 0.5% blocking reagent (Roche), followed by treatment with 20% heat-inactivated sheep serum (Sigma-Aldrich). After incubation with an alkaline phosphatase (AP)-conjugated anti-DIG antibody (Roche) for 2 h, the sections were visualized using a NBT/BCIP solution (Roche).

### Antibodies

The hybridoma cells producing anti-TMEM180 monoclonal antibodies (mAbs) were established using myeloma cells (p3×63) and lymph-node cells from a rat with immunizing recombinant human extradomain (355–400 aa, UniProtKB entry number Q14CX5) of TMEM180 or TMEM180-positive tumour exosomes. The TMEM180-positive tumour exosomes were purified from the culture supernatant of DLD-1-OE cells using hydroxyapatite and gel filtration (Superdex200pg, GE healthcare). Initially, the anti-TMEM180 mAb (rat IgM) was obtained using recombinant human extradomain of TMEM180. The expression level of TMEM180 in the exosome was confirmed by an ELISA test with the mAb. Finally, the anti-TMEM180 mAb (rat IgM, clone 669) was obtained using TMEM180-positive tumour exosomes. The heavy-chain variable and the kappa light-chain variable-region cDNAs were humanized and cloned into the vector for the human IgG1 expression. The vectors were transfected into CHO cells (Riken BioResource Center). A stable clone (humanized IgG, clone 669) was isolated. The procedure was conducted according to standard protocols^42^.

### Exosome isolation, immunogold electron microscopy and ELISA

Here, 6.8 × 10^6^ DLD-1 cells were plated on a 15-cm dish and grown overnight. The FBS-supplemented medium was depleted by washing twice with PBS. The cells were then incubated with serum-free culture medium at 30 ml/dish for 24 h. After the collection and filtration through a 0.22-µm filter, the exosome-containing supernatant was obtained. This supernatant was preserved at 4°C with a protease inhibitor (Wako) in each experiment. For the immunogold electron microscopy, isolated exosomes were adsorbed on nickel grids with supported films. The anti-TMEM180 mAb, anti-CD9 (Ancell) or CD63 mAb (Ancell) was added into the grids, which were then incubated at room temperature. After the grids were washed with 1% BSA/PBS, they were incubated at room temperature with the gold-conjugated goat anti-rat/mouse IgG polyclonal antibody (British BioCell International Ltd.) and washed with 1% BSA/PBS. After the glutaraldehyde fixation, negative staining was performed using a 2% solution of phosphotungstic acid (pH 7.0). The samples were then dried and visualized with transmission electron microscopy (JEM-1400Plus, JEOL Ltd.). Images were acquired with a CCD camera (VELETA, Olympus Soft Imaging Solutions GmbH).

For the sandwich ELISA test, anti-TMEM 180 mAb or anti-CD9 mAb (Ancell) as a capture antibody was coated onto 96-well plates. Isolated exosomes form DLD1 or K562 cells were added into the each palate and incubated for 1 h at room temperature. After the washing, the plates were followed by incubation with HRP-labelled anti-TMEM180 mAb as a detection antibody for 1 h at room temperature. After the washing, the absorbance at 450nm was measured by Spectra Max paradigm (Molecular Devices).

### Immunohistochemistry

Human specimens were purchased from the BioChain Institute, Inc.. Immunohistochemistry was performed as previously described method^43^. Anti-TMEM180 antibody or anti-EGFR mAb (cetuximab, Merck) was labelled with HRP using Peroxidase Labeling Kit-SH (Dojindo). The tissues were then incubated with the HRP-conjugated anti-TMEM180 mAb or anti-EGFR mAb for 1 h at room temperature. After the tissue was washed with PBS, the reaction was visualized using DAB (Dako), and the slides were counterstained with haematoxylin. Frozen tumour samples were obtained from the DLD-1 and SW480 xenograft mouse models. They were fixed with 4% PFA in PBS for 20 min at room temperature and blocked with 5% skim milk (Becton Dickinson) in PBS for 1 h at room temperature. Then sections were sequentially stained with Rabbit anti-alpha SMA antibody (1:100, abcam) for 1h at room temperature, and then with goat anti-rabbit IgG (H+L) Alexa Fluor 488 (1:100, Thermo fisher Scientific) for 1h at room temperature. Sections were stained with rat anti-integrin alpha 6 antibody (1:100, abcam) for 1h at room temperature, and then with goat anti-rat IgG (H+L) Alexa Fluor 555 (1:100, Thermo fisher Scientific). Slices were also stained with anti-TMEM180 antibody conjugated with HRP (1:100), for 1h at room temperature, and then with anti-HRP antibody Alexa Fluor 647 (1:250, Jackson ImmunoResearch) for 1 h at room temperature in sequence. The slides were washed and counterstained with DAPI solution (Thermo Fisher Scientific). Each image was acquired using the VS120 Virtual Microscope System (Olympus). The data were analysed using the OlyVIA program (Olympus).

For the cell staining, cultured cells were fixed with 4% PFA/PBS. After the permeabilization with 0.1% Triton X-100/PBS and blocking with 5% skimmed milk, the cells were stained with R-phycoerythrin-labelled anti-TMEM180 using R-Phycoerythrin Labeling Kit-SH (Dojindo) as the primary antibody. The cells were then incubated with the goat anti-mouse IgG antibody conjugated to Alexa Fluor 488 (1:200, Thermo Fisher Scientific) as a secondary antibody and DAPI solution for nuclear staining. Images were acquired using a BZ-9000 digital high-definition microscopic system (Keyence Co.). The fluorescence intensity of TMEM180 from at least 1500 DAPI positive cells were analysed using ImageJ (NIH). Representative results were from at least two independent experiments.

### Hematoxylin and eosin staining

The sections were fixed with 4% PFA in PBS for 20 min at room temperature. Then the sections were stained with Mayer’s Hematoxylin Solution (Muto Pure Chemicals) for 10 min at room temperature and Eosin solution (Muto Pure Chemicals) for 3 min at room temperature.

### Western blot analysis

The cells were lysed with buffer containing 20 mM Tris-HCl (pH 7.5), 150 mM NaCl, 0.1% sodium dodecyl sulfate (SDS), 1% Triton-X100 and cOmplete protease inhibitor cocktail (Roche). The lysate was boiled at 95°C for 5 min for denaturing and inactivation of the endogenous proteases. The samples were separated using SDS-Gel electrophoresis (4– 15% Mini-PROTEAN TGX precast gels, Bio-Rad) and transferred onto polyvinyldifluoride membranes (Bio-Rad). The membranes were blocked with tris-buffered saline (TBS) containing 5% skimmed milk and 0.1% Tween 20 (Sigma-Aldrich). The blots were then incubated with anti-HIF-1α (1:500, GeneTex) or β-Actin (1:1000, Epitomics) antibodies as the primary antibody at room temperature for 2 h or at 4°C overnight. The membranes were washed three times with TBS containing 0.1% Tween 20 (TBS-T) and then incubated with HRP-conjugated anti-mouse (1:1000, R&D Systems) or anti-rabbit IgG (1:1000, CST) as the secondary antibody. The Can Get Signal solution (Toyobo) was used to reduce the background noise. After the membranes were washed with TBS-T, the proteins were visualized using ECL prime (GE Healthcare). The images were acquired using a ChemiDoc imager (Bio-Rad).

### FACS analysis

FACS analysis and cell sorting were conducted as previously described method^9^. *In vitro* cultured cells were stained with anti-TMEM180 mAbs (clone 669) and anti-CD44 mAb (NeoMarkers) as a first antibody, and an Alexa Fluor 488/647-conjugated anti-mouse/human-Ig polyclonal antibody (Thermo Fisher Scientific) as a secondary antibody. The stained cells were analysed using a Guava easyCyte 10HT (Merck Millipore Co.) or Aria flow cytometer (BD Biosciences). The dead cells, which were stained using propidium iodide (PI) (Thermo Fisher Scientific), were excluded from the analysis. The data were analysed using the FlowJo program (Tree Star).

### ADCC and CDC assay

^51^Cr release assays were conducted to evaluate the ADCC and CDC activities. Here, 5 × 10^6^ DLD-1, SW480 or K562 cells were labelled for 2 h at 37°C with 3.7 MBq of ^51^Cr (100 μCi).

For the ADCC evaluation, cells were placed in 96-well plates at 1 × 10^5^ or 1 × 10^4^ cells/well. The cells were exposed to the anti-TMEM180 mAb or anti-EGFR mAb at the 40 fold times number of NK cells (Chemicals Evaluation and Research Institute) as an effector target ratio and used as effector cells in this assay.

For the CDC evaluation, cells were placed in 96-well plates at 5 × 10^4^ cells/well. The cells were exposed to the anti-TMEM180 mAb or anti-EGFR mAb with complement human sera (Chemicals Evaluation and Research Institute).

The cells were then incubated at 37°C, 5% CO_2_ for 4 h. Cytotoxicity (%) = [(test sample release-background release)/(maximum release - background release)] × 100. All samples were run in triplicate.

### Animal model, tumour-initiating activity and anti-tumour activity

Female BALB/c nude mice (5 weeks old) were purchased from Charles River Laboratories Japan Inc. For the evaluation of tumour-initiating activity, SW480-WT/-KD cells were suspended in 100 μl of PBS. The mice were inoculated subcutaneously in the left flank with 1 × 10^6^ or 1 × 10^5^ cells.

To evaluate the anti-tumour activity of the anti-TMEM180 mAb, mice were inoculated subcutaneously in the flank with 5 × 10^6^ DLD-1 or SW480 cells. The length (L) and width (W) of tumour masses were measured every four days, and tumour volume was calculated using (L×W2)/2. When the mean tumour volume reached approximately 100–200 mm^3^, the mice were randomly divided into groups that consisted of five mice each. PBS, the anti-TMEM180 mAb or the anti-EGFR mAb cetuximab was administered on day 0 by intraperitoneal injection.

All animal procedures were performed in compliance with the Guidelines for the Care and Use of Experimental Animals that were established by the Committee for Animal Experimentation of the National Cancer Center. These guidelines meet the ethical standards required by law and comply with the guidelines for the use of experimental animals in Japan.

### Database analysis

Hypoxia response element consensus sequences (ACGTG or GCGTG)^12,13^ at 3,000 bp upstream from exon 1 of the *TMEM180* gene were analysed using NCBI Gene databases (https://www.ncbi.nlm.nih.gov/gene/). *TMEM180* gene expression in normal tissues and cancer tissues was analysed using the microarray gene expression dataset from RefEx (http://refex.dbcls.jp/)^44^ and Oncomine (http://www.oncomine.org/)^45^, respectively. The comparison of *TMEM180* gene expression between 286 colorectal adenocarcinoma patient samples and 41 normal tissue samples using TCGA cancer genomics data was performed using UALCAN (http://ulcan.path.uab.edu/index.html) ^46^.

### Generation of TMEM180 KO mice

TMEM180 KO mice were generated using CRISPR/Cas9-mediated genome editing technology^47^. Briefly, guide RNAs (gRNAs) at exon 2 of mouse *TMEM180* genome immediately downstream of the ATG start codon were designed using CRISPRdirect^48^. Two gRNAs [gS01(5’-AGCTGTGGTGTACGGCTCGTTGG-3’), gS03(5’-TGTGTTCCTGCTGTACTACGTGG-3’)] were selected *in vitro* digestion assay system^49^ using the target *TMEM180* PCR product. These 2 gRNAs were inserted into the *Bbs*I restriction site in the px330 plasmid and constructed as *px330-TMEM180* genome editing vector.

The pronuclear stage ova of C57BL/6J (CLEA) were prepared by in vitro fertilization and injected 1.67 ng/ul of *px330-TMEM180* DNA vector into the pronucleus according to standard protocols^50^. The injected ova were transferred into the oviduct of the pseudopregnant Jcl:MCH (CLEA) female on that day.

Two viable mice from gS01 and 14 viable mice from gS03 were obtained independently via caesarean section. Genotyping of mouse tail DNA via PCR [Forward (5’-CTAATAGGCAACCGCAGAGC-3’), Reverse (5’-CTAAACAGCACAGTCTGCCC-3’)] and purified PCR products were sequenced using the forward primer. The results of the sequence analysis are shown in Supplemental Table1.

### Statistical analysis

Significant differences between the groups were determined using Student’s t-test for ELISA data (Fig. 3c, e, Fig. 4c and Supplementary Fig. 1c), ANOVA to compare the tumour size (Fig. 2a) and body weight (Fig. 2b), and chi-squared test (Supplementary Fig. 3b). All analyses were performed using SPSS software version 20 (IBM).

---

**Supplementary Figure 1.**
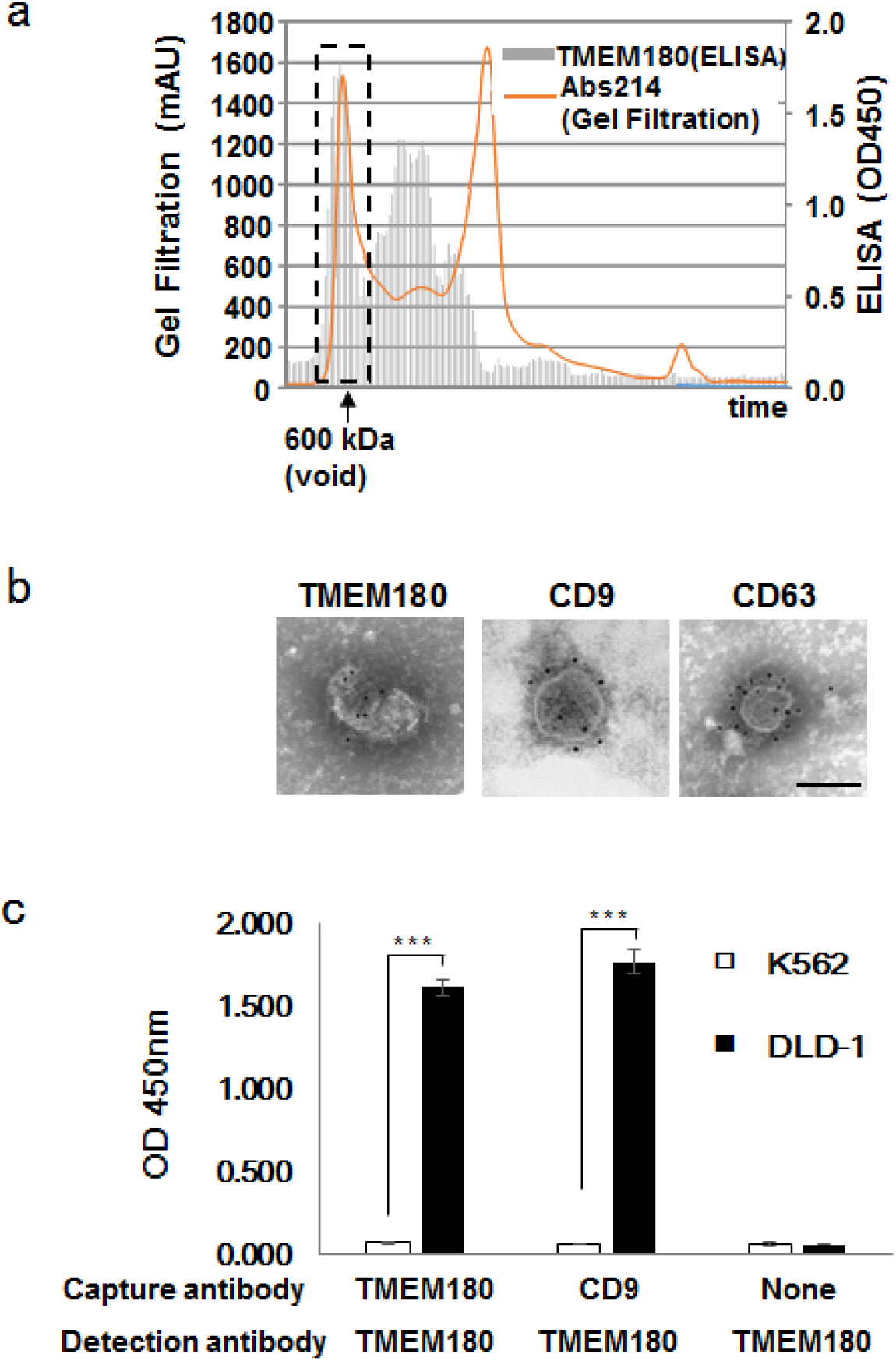
TMEM180 expression in tumour exosomes. a. Purification of the TMEM180 protein from DLD-1 cells for the immunization. Gel filtration chromatography showed that a high concentration of TMEM180 protein existed in the void fraction with a molecular weight over 600 kDa. b. Immunogold electron microscopy. TMEM180 expression in tumour exosomes was determined. CD9 and CD63 were used as positive controls of standard exosome markers. Scale bar = 100 nm. c. Sandwich ELISA. Both TMEM180-positive and CD9 positive exosomes derived from DLD1 cells were determined. K562 cells were used as negative control. ***P < 0.001. Bar = SD.

**Supplementary Figure 2.**
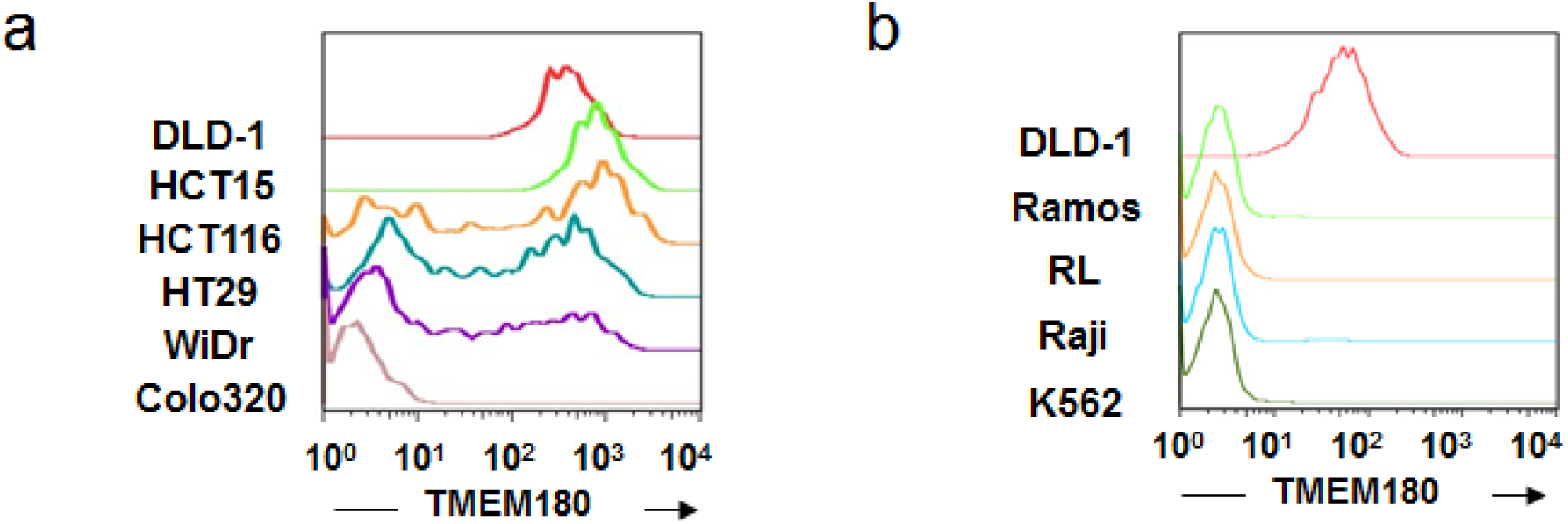
TMEM180 expression in CRC and hematopoietic cell lines. a and b. Flow cytometric analysis of the colorectal cancer (CRC) cell lines DLD-1, HCT15, HCT116, HT29, WiDr and Colo320 (a). Flow cytometric analysis of haematopoietic cell lines Ramos, RL, Raji and K562 (b). DLD-1 was used as a positive control for TMEM180 expression.

**Supplementary Figure 3.**
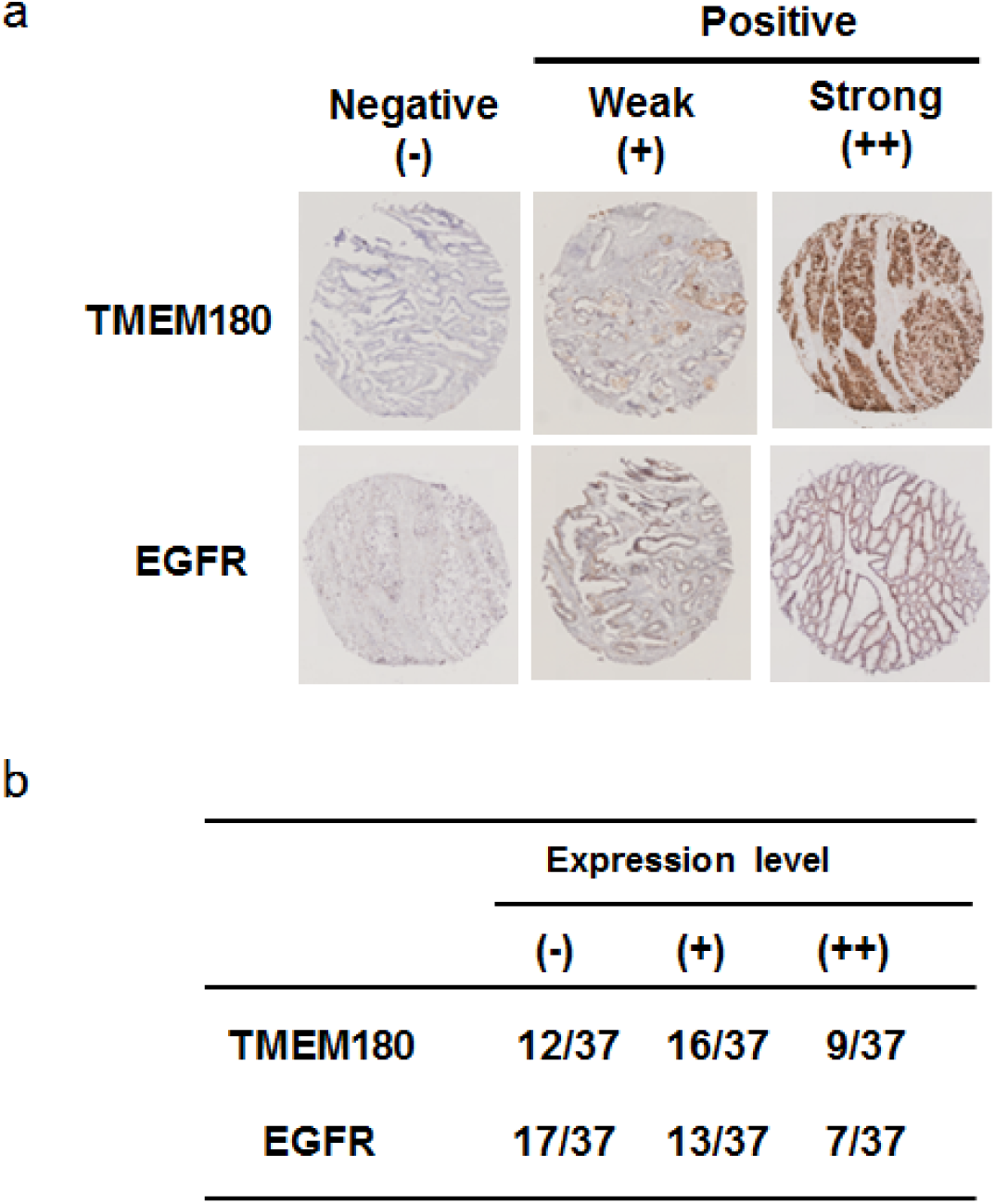
TMEM180 and EGFR expression level in CRC tissues. a. Representative immunostaining patterns of TMEM180 and EGFR in CRC tissues were classified as negative (-), a weak positive (+) and strong positive (++). b. The number of cases with each expression level (-), (+) or (++) of TMEM180 or EGFR in CRC tissues (total number 37) was shown.

**Supplementary Figure 4.**
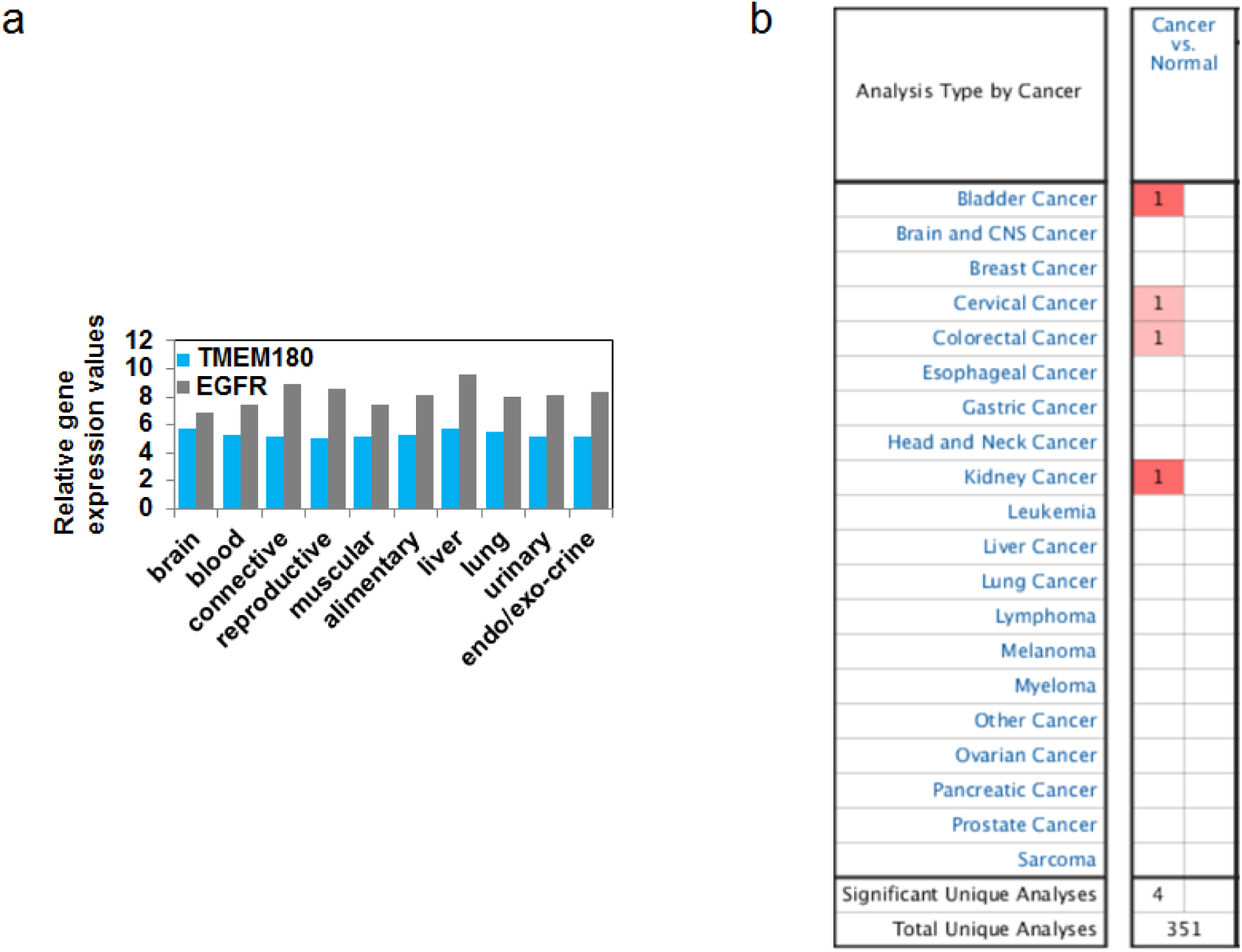
TMEM180 expression in cancer and normal tissues. a. Representative immunostaining patterns of TMEM180 and EGFR in CRC tissues were classified as negative (-), a weak positive (+) and strong positive (++). b. The number of cases with each expression level (-), (+) or (++) of TMEM180 or EGFR in CRC tissues (total number 37) was shown.

**Supplementary Figure 5.**
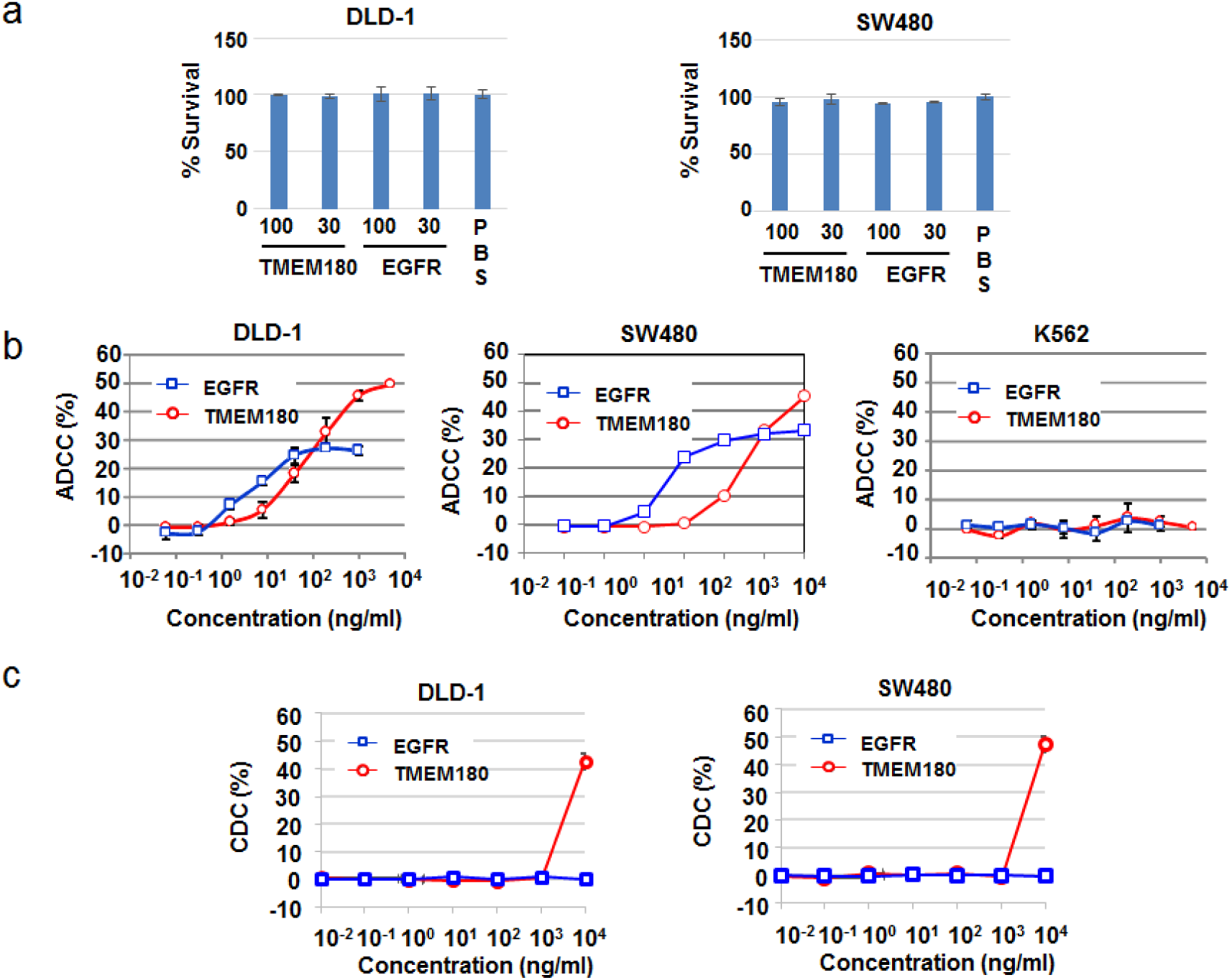
*In vitro* cytotoxicity and ADCC or CDC activity of the anti-TMEM180 mAb. a. *In vitro* cytotoxicity of the anti-TMEM180 mAb for CRC DLD-1 or SW480 cells in the WST-8 assay. The anti-TMEM180 mAb (=TMEM180), the anti-EGFR mAb cetuximab (=EGFR) or saline (=PBS) as a control was given at a concentration of 30 or 100 μg/ml. The percentage of surviving cells was calculated as the ratio of the number of cells in each group to that in the group given PBS treatment. b and c. ADCC (antibody-dependent cellular cytotoxicity, b) or CDC (complement-dependent cytotoxicity, c) activity of the anti-TMEM180 mAb for CRC DLD-1 cells, CRC SW480 cells and haematopoietic K562 cells. Red and blue lines indicate the ADCC or CDC activity of the anti-TMEM180 mAb (=TMEM180) and anti-EGFR mAb cetuximab (=EGFR), respectively. Bar = SD.

**Supplementary Figure 6.**
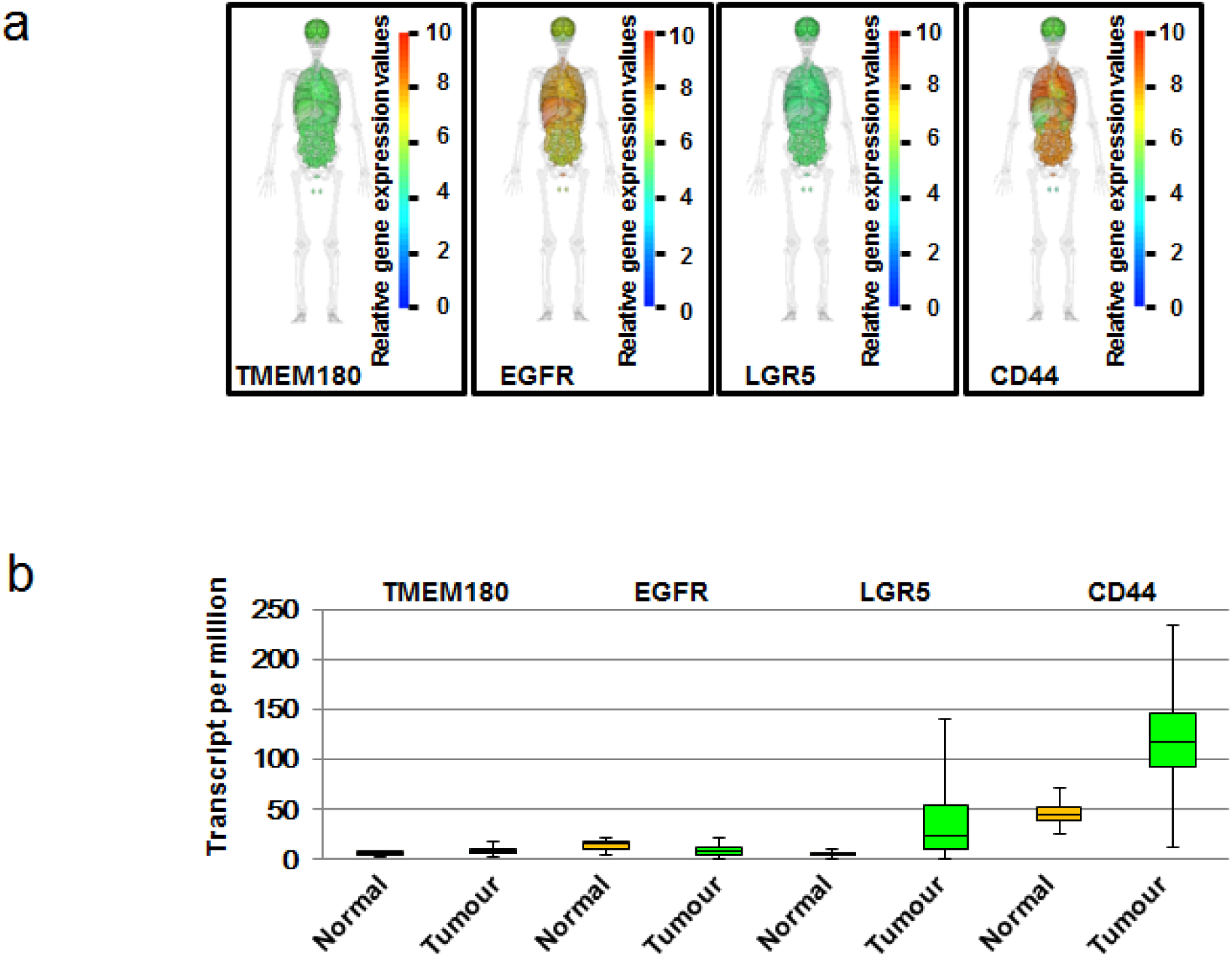
*TMEM180* mRNA expression compared with *EGFR, LGR5* and *CD44* expression. *a. TMEM180, EGFR, LGR5* and *CD44* mRNA expression in ten normal tissues (brain, blood, connective, reproductive, muscular, alimentary, liver, lung, urinary and endo/exocrine tissue). Dataset was obtained from RefEx (http://refex.dbcls.jp/). b. Comparison of *TMEM180, EGFR, LGR5* and *CD44* mRNA expression between colorectal cancer (N = 286) and normal tissue (N = 41). Dataset was obtained from UALCAN (http://ulcan.path.uab.edu/index.html).

**Supplementary Table 1:**
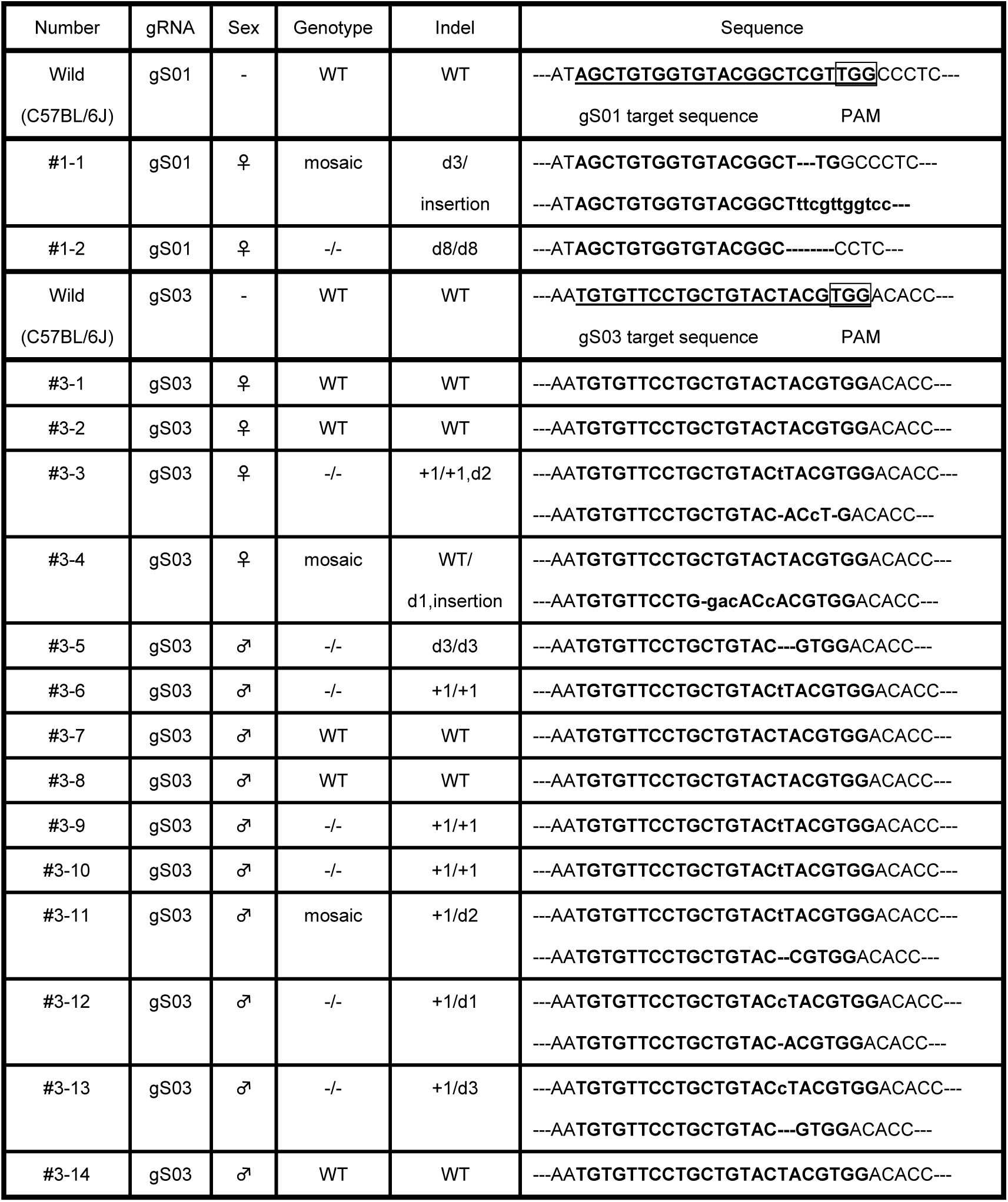
**Genotyping analysis of TMEM180 KO F0 mice**

## Acknowledgements

This work was supported in part by a research and development by the New Energy and Industrial Technology Development Organization (NEDO), National Cancer Center Research and Development Fund (26-A-14, 26-A-12, 29-A-9, 29-S-1), and Project for Cancer Research and Therapeutic Evolution from the Japan Agency for Medical Research and Development (AMED) (17cm0106415h0002).

## Author Contributions

M.Y and Y.M. designed the experiments and identified TMEM180. S.S., S.H. and T.A. established the monoclonal antibody and produced humanized antibody. R.T. and S.S. performed the experiments of CSC *in vitro* and *in vivo*. T.A. performed the experiments of HIF-1 and data base analysis. M.Y., S.S., S.H., T.A., R.T. and Y.M. wrote the manuscript. Y.M. conceived and supervised the project.

